# Dynamic Formation of the Protein-Lipid Pre-fusion Complex

**DOI:** 10.1101/2024.04.17.589983

**Authors:** Maria Bykhovskaia

## Abstract

Synaptic vesicles (SVs) fuse with the presynaptic membrane (PM) to release neuronal transmitters. The SV protein Synaptotagmin 1 (Syt1) serves as a Ca^2+^ sensor for evoked fusion. Syt1 is thought to trigger fusion by penetrating into PM upon Ca^2+^ binding, however the mechanistic detail of this process is still debated. Syt1 interacts with the SNARE complex, a coiled-coil four-helical bundle that enables the SV-PM attachment. The SNARE-associated protein Complexin (Cpx) promotes the Ca^2+^-dependent fusion, possibly interacting with Syt1. We employed all-atom molecular dynamics (MD) to investigate the formation of the Syt1-SNARE-Cpx complex interacting with the lipid bilayers of PM and SV. Our simulations demonstrated that the PM-Syt1-SNARE-Cpx complex can transition to a “dead-end” state, wherein Syt1 attaches tightly to PM but does not immerse into it, as opposed to a pre-fusion state, which has the tips of the Ca^2+^-bound C2 domains of Syt1 inserted into PM. Our simulations unraveled the sequence of Syt1 conformational transitions, including the simultaneous Syt1 docking to the SNARE-Cpx bundle and PM, followed by the Ca^2+^ chelation and the penetration of the tips of Syt1 domains into PM, leading to the pre-fusion state of the protein-lipid complex. Importantly, we found that the direct Syt1-Cpx interactions are required to promote these transitions. Thus, we developed the all-atom dynamic model of the conformational transitions that lead to the formation of the pre-fusion PM-Syt1-SNARE-Cpx complex. Our simulations also revealed an alternative “dead-end” state of the protein-lipid complex that can be formed if this pathway is disrupted.

**Statement of Significance:** Neurons communicate by releasing transmitter molecules. Transmitters are packed in synaptic vesicles (SVs) and released by the fusion of SVs with the presynaptic membrane (PM). This process is regulated by a dynamic complex of fusion proteins, including the coil-coiled SNARE bundle that attaches SV to PM, Synaptotagmin that serves as a Ca^2+^ sensor for the release process, and Complexin that attaches to the SNARE bundle and promotes the Ca^2+^-dependent release. To understand how these proteins interact dynamically with the lipid bilayers of SV and PM, we employed molecular dynamics, a computational approach that enables simulating the behavior of proteins and lipids at the atomistic resolution. Our simulations enabled us to delineate the stages of the formation of prefusion protein-lipid complex.

## Introduction

Neuronal transmitters are packaged in synaptic vesicles (SVs) and released by the fusion of SVs with the presynaptic membrane (PM). Action potentials trigger SV-PM fusion by inducing an inflow of Ca^2+^ into the nerve terminal, and the SV protein Synaptotagmin 1 (Syt1) serves as a Ca^2+^ sensor [3, 4]. Syt1 is a tansmembrane (TM) protein with its cytosolic region being comprised of two domains, C2A and C2B [5]. Each domain has two loops forming a Ca^2+^ binding pocket, and in each of the pockets Ca^2+^ ions are chelated by five aspartic acids [6, 7]. It is agreed that the synergistic coordinated insertion of the tips of the C2 domains into the phospholipid membrane drives fusion [3, 8, 9], but other mechanistic details of Syt1 action are still debated.

Syt1 is interacts with the SNARE complex[10–12], a coil-coiled four-stranded helical bundle made up of the PM-associated proteins (also known as t-SNARE) syntaxin-1A (Sx) and SNAP25, as well as the SV protein synaptobrevin (Sb) or v-SNARE. The SNARE complex mediates the SV-PM attachment, and the SNARE assembly overcomes the electrostatic and hydration repulsion between the lipid bilayers [13]. Multiple studies suggest an important role for the Syt1-SNARE interactions [1, 14–18], however it is still debated how the Syt1-SNARE complex is formed *in vivo*. A cytosolic protein Complexin (Cpx) attaches to the SNARE bundle[19] and serves as a positive regulator of synchronous release, promoting and accelerating evoked synaptic transmission[20–25]. The effect of Cpx on synchronous fusion is Ca^2+^-dependent [20] and several studies suggested a functional [25–29] or molecular [30] interaction between Cpx and Syt1. However, it is still debated how Cpx promotes and synchronizes evoked transmission.

The Syt1-SNARE complex adopts multiple conformational states[1, 18, 30–32], and it is still obscure how the complex is formed between the lipid bilayers of SV and PM. The molecular dynamics (MD) approach proved to be instrumental in addressing this problem [33, 34], however, the simulation time-scales represent a critical limitation of MD. Here, we took advantage of Anton2 supercomputer customized for MD [35, 36], which enables the simulations of protein-lipid complexes at a time-scale of microseconds and up a sub-millisecond range [37, 38]. We have recently employed this approach to build the all-atom model of the PM-Syt1-SNARE-Cpx complex [2]. The present study investigates the conformational pathway of the complex leading to its prefusion state.

## Methods

### System setup

All the molecular systems were constructed using Visual Molecular Dynamics Software (VMD, Theoretical and Computational Biophysics Group, NIH Center for Macromolecular Modeling and Bioinformatics, at the Beckman Institute, University of Illinois at Urbana-Champaign). All the simulations were performed in water/ion environment with explicit water molecules. Potassium and chloride ions were added to neutralize the systems and to yield 150 mM concentration of KCl.

The lipid bilayer was constructed as described in [2].The phosphatidylcholine (POPC) lipid bilayers were generated using VMD. To mimic PM, the anionic lipid patch containing POPC, phosphatidylserine (POPS) and PIP_2_ (75:20:5) was used. The initial structure of the POPC:POPS:PIP_2_ bilayer [39] was kindly provided by Dr. J. Wereszczynski (Illinois Institute of Technology). We have manually generated the bilayer with one leaflet containing only POPC monomers, and the other leaflet having the POPC:POPS:PIP_2_ (75:20:5) composition. The leaflets were merged using VMD and then optimized and equilibrated in water/ion environment. This approach ensured that the periodic image of the lipid bilayer mimicking PM served to mimic the neutral lipids composing SV. In all the systems, the lipid bilayers were positioned in the XY plain.

The model of full length Syt1^fl^ was generated employing the SWISS-MODEL server [40, 41].The sequence of human Syt1 (GeneBank: KAI2567145.1), excluding its luminal domain (residues 1-55) was used [42], and the structure of the C2AB tandem of Syt1[43] served as a template for generating the 3D structure of Syt1^fl^. The model of the SNARE-Cpx complex was taken from earlier MD simulations [2, 44].

### Molecular dynamics

The MD simulations were performed employing CHARMM36 force field [45] modified to include the parameters for PIP_2_ as described in [39]. All the simulations were performed with periodic boundary conditions and Ewald electrostatics at 310K.

The heating (20 ps) and equilibration (100 ns) phases were performed employing NAMD [46] Scalable Molecular Dynamics (Theoretical and Computational Biophysics Group, NIH Center for Macromolecular Modeling and Bioinformatics, at the Beckman Institute, University of Illinois at Urbana-Champaign) through ACCESS (Advanced Cyberinfrastructure Coordination Ecosystem: Service and Support) at Stampede (TACC) and Anvil (Purdue) clusters. The NAMD simulations were performed in NPT ensemble with a flexible cell and with a time-step of 1.5 fs, employing Langevin thermostat and Berendsen barostat.

Production runs were performed at Anton2 supercomputer [35, 36] with Desmond software through the MMBioS (National Center for Multiscale Modeling of Biological Systems, Pittsburgh Supercomputing Center and D.E. Shaw Research Institute). All the Anton2 simulations were performed in a semi-isotropic regime, with a time-step of 2.5 fs, employing the multigrator [47] to maintain constant temperature and pressure.

All the trajectories are summarized in the Table 1.

**Table 1.**
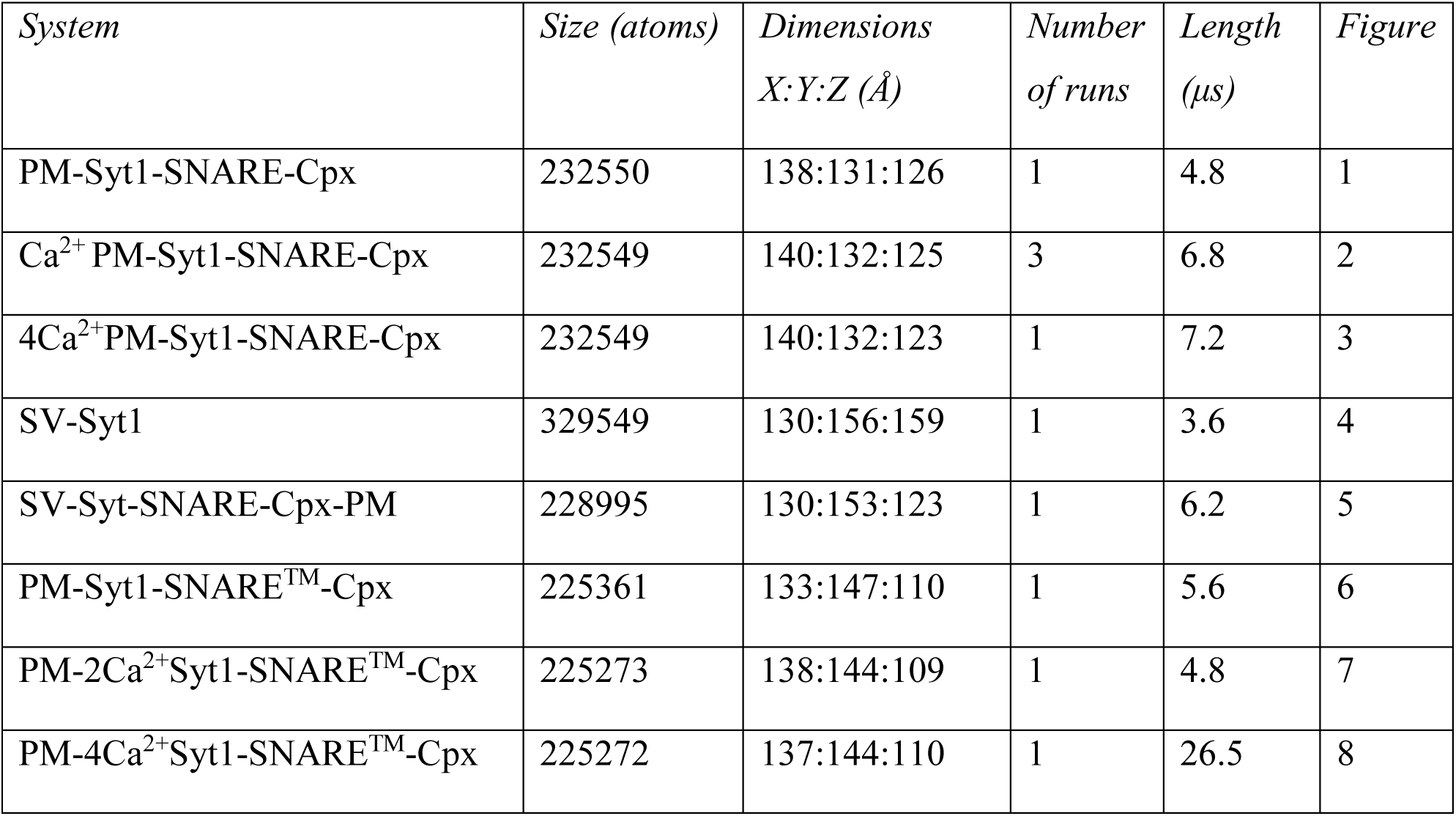
MD trajectories.

### Analysis and Visualization

The trajectory analysis was performed employing VMD. All the parameters along all the trajectories were computed with a time step of 2.4 ns. The number of VdW contacts between two molecules was computed as the number of atomic pairs separated by less than 3 Å.

The penetration of the CBLs of the Syt1 domains into the lipid bilayer of PM (CBL-PM) was computed as 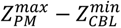, where 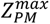 is the maximal atomic Z coordinate of the lipid bilayer, and 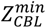 is the minimal atomic Z coordinate of the CBL motif of either C2A or C2B domain of Syt1. Note that the negative values of the parameter (CBL-PM) correspond to CBLs being separated from PM, while the values of 1 nm or higher correspond the penetration of CBLs into PM[2].

Each trajectory was analyzed visually to ensure that no element of the system came in contact with an element of the periodic image. The only exception was the system SV-Syt-SNARE-Cpx-PM (Table 1, Fig. 5), which had by design Syt1 and SNARE proteins contacting PM represented as the periodic image of SV.

The distributions of the parameter along MD trajectories were compared using the Kolomogorov-Smirnov (K.-S.) test.

## Results

### The PM-Syt1-SNARE-Cpx complex can adopt the “dead-end” state, which has Syt1 tightly attached to PM but not penetrating into the lipid bilayer

The primary interface between t-SNARE and the C2B domain of Syt1 (strands β1, β2, β5, and β8) has been identified by crystallography[1]. A subsequent *in vivo* study in *Drosophila* revealed multiple loss-of-function mutations in Syt1 pertaining to this interface[17], and MD simulations demonstrated that its formation promotes the immersion of the tips of the C2 domains of Syt1 into PM and drives the SV-PM fusion [2]. However, multiple conformational states of the Syt1-SNARE complex have been detected *in vitro*[18, 31] and *in silico*[2], raising the question of whether some of these states can serve as intermediates in the formation of the pre-fusion protein-lipid complex. To address this question, we employed the MD approach to test two scenarios: 1) Syt1 anchors to the SNARE bundle via the primary interface; and 2) Syt1 anchors to the SNARE bundle forming an intermediate state and subsequently undergoes conformational transition(s) leading to the pre-fusion state.

As a starting point, we took the Syt1-SNARE-Cpx complex obtained by crystallography [1] followed by MD [2], which had the C2B domain of Syt1 attached to t-SNARE via its primary interface. The protein complex was then positioned onto the lipid bilayer mimicking PM (Fig. 1 A), so that the PM-binding motifs of Syt1, including its polybasic (PB) region [48, 49] and Ca^2+^-binding loops (CBLs)[3] contacted PM. A subsequent MD run (4.8 µs, Video 1, Fig. 1 B) produced a conformational transition, so that the C2A domain separated from the SNARE bundle, attached to its N-terminus (Fig. 1 C, Transitional state) and then attached to PM (Fig. 1 C, Final state). The final 3 µs of the simulations (Fig. 1 B) did not produce any further alterations: both C2 domains remained attached to PM with their CBLs facing each other (Fig. 1 D).

**Figure. 1.**
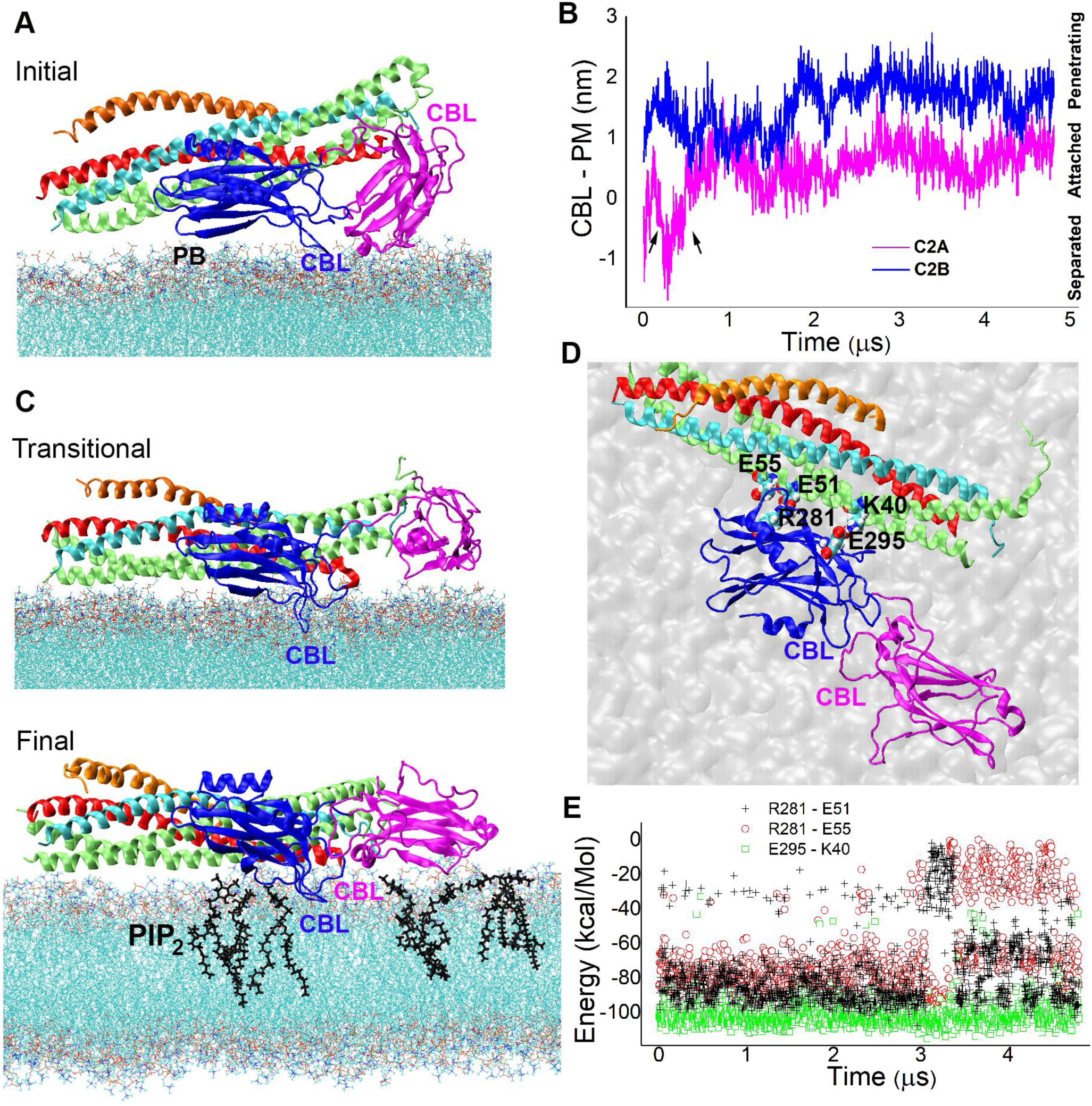
The attachment of Syt1 to the SNARE-Cpx complex via the primary interface produces the PM-Syt1-SNARE-Cpx complex with the CBLs of the C2 domains facing each other. **A.** The Syt1-SNARE-Cpx complex with the C2B domain attached to t-SNARE via the primary interface[1, 2] is positioned onto PM in such a way that PB and CBL of the C2B domain are in contact with PM. Magenta: C2A domain; blue: C2B domain; red: Sb; cyan: Sx; lime: SNAP25; orange: Cpx. **B.** The C2A domain undergoes a conformational transition (arrows) and then attaches to PM. The parameter CBL-PM was computed as described in Methods. **C.** The transitional and final states of the PM-Syt1-SNARE-Cpx complex. Note that the final state has both domains anchored to PM via interactions with PIP_2_ (black). **D.** The top view of the final PM-Syt1-SNARE-Cpx complex showing the primary C2B-SNARE interface maintained by the salt bridges between the C2B domain of Syt1 and SNAP25 (shown in VdW representation). Note the CBLs of the C2 domains of Syt1 facing each other. **E.** The salt bridges maintaining the CBL-SNARE interface along the trajectory.

The attachment of both C2 domains to PM was maintained by salt bridges of the basic residues of Syt1 with phosphotidylinositon 4,5-biphosphate (PIP_2_) monomers of PM (Fig. 1 B, black). The C2B-SNARE interface remained intact, being largely maintained by the salt bridges between R281 of Syt1 and E51/E55 of SNAP25, as well as between E295 of Syt1 and K40 of SNAP25 (Fig. 1 D, E).

Next, we investigated how Syt1 binds Ca^2+^ within the obtained PM-Syt1-SNARE-Cpx complex. Ca^2+^ chelation was simulated employing the method of K^+^/Ca^2+^ replacement[50]. We examined the final 250 ns stretch of the MD trajectory (presented in Fig. 1) and identified all the time-points which had K^+^ ions inside the Ca^2+^ binding pockets of both C2 domains. We then selected three time-points which had three K^+^ ions inserted between the CBLs of the C2A domain[6] and two K^+^ ions inserted between the CBLs of the C2B domain [51]. We next replaced the five K^+^ ions by Ca^2+^ and performed three 6.8 μs MD runs starting from the three initial states described above.

Surprisingly, all the three MD runs produced the complexes with Ca^2+^ ions bridging the CBLs of the C2A and C2B domains (Fig. 2). In each run, one or two Ca^2+^ ions dissociated from Syt1, leaving three or four Ca^2+^ ions chelated. The first run (Run 1) had four Ca^2+^ ions chelated in the end of the trajectory (Fig. 2 A), with two Ca^2+^ ions bridging the C2A and C2B domains, and each of the other two Ca^2+^ ions being chelated by either C2A or C2B domain only. The fifth Ca^2+^ ion dissociated from Syt1 after 3 μs of the simulation (Fig. 2 B, note one of the ions having zero interaction energy with Syt1, ochre). The other two runs (Run 2 and 3) had three Ca^2+^ ions remaining chelated in the end of the trajectory, with one of the chelated Ca^2+^ ions bringing the CBLs of the C2A and C2B domains (Fig. 2 C).

**Figure 2.**
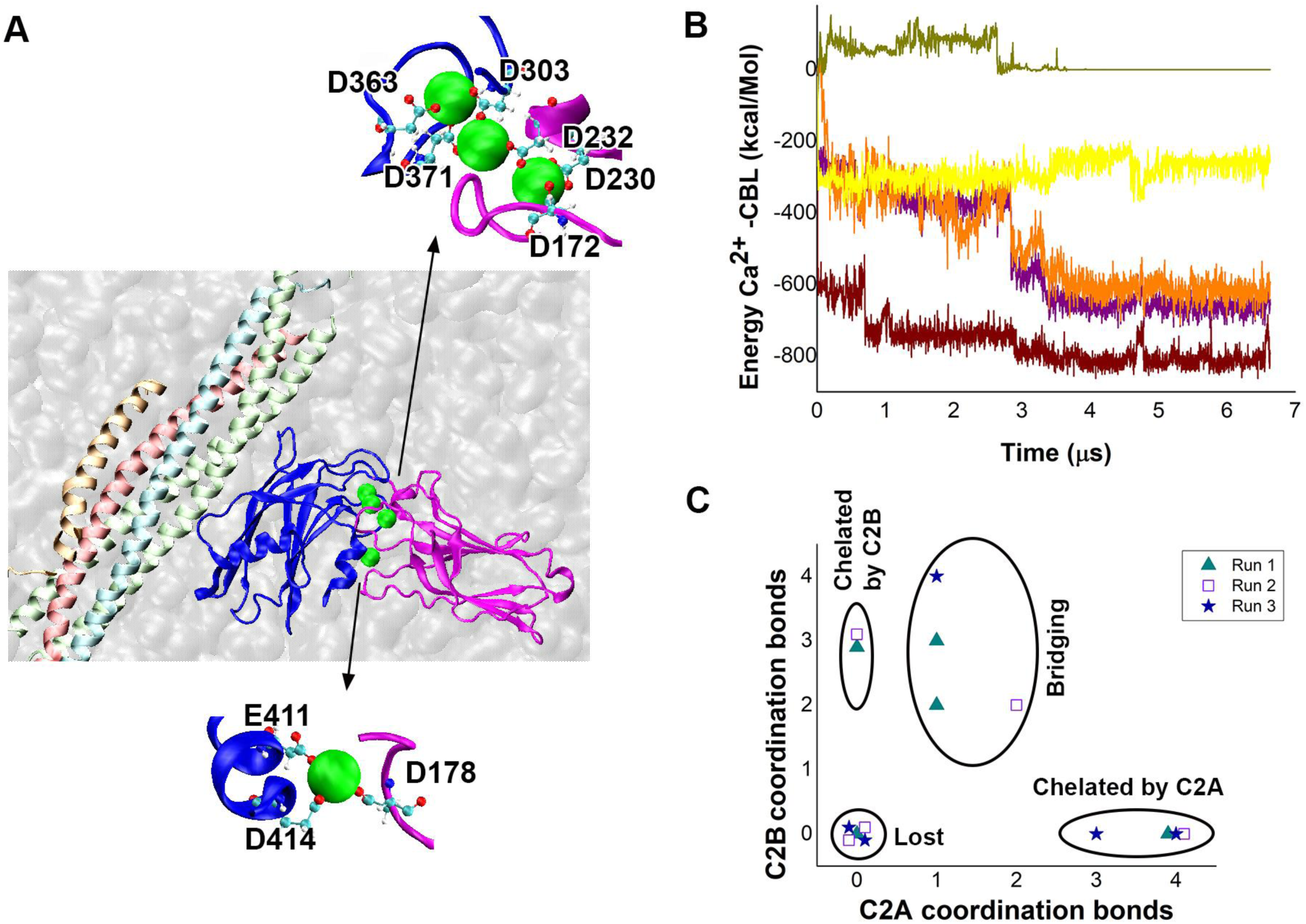
Ca^2+^ ions chelated within the PM-Syt1-SNARE-Cpx complex bridge the CBLs of the C2A and C2B domains. **A.** The PM-4Ca^2+^Syt1-SNARE-Cpx complex in the end of the Run 1. Four Ca^2+^ ions (green spheres) are chelated. One of the ions is bridging the CBLs of the C2 domains (D303, D371, D232) while another ion is bridging the CBL of the C2A domain (D178) with the C-terminus of the C2B domain (E411 and E414). **B.** The energy of the interactions of the five Ca^2+^ ions (marked by different colors) with the CBLs of Syt1 over the course of the Run 1. Note that one of the ions (ochre) is “lost” after the initial 3 µs, as the interaction energy approaches zero. The remaining four Ca^2+^ ions formed stable coordination bonds. **C.** The results of the three MD runs. Each data point corresponds to a single Ca^2+^ ion in the end of the trajectory. Note that each run produced bridging of the C2A-C2B domains by either one (Runs 2 and 3) or two (Run 1) Ca^2+^ ions.

These results show that the proximity to PM can drastically alter the coordination of Ca^2+^ ions chelated by Syt1. We show that Syt1 attached to PM binds Ca^2+^ in such a way that Ca^2+^ ions bridge C2 domains. In contrast, it was demonstrated *in vitro* [1] and *in silico* [50] that when phospholipid bilayers are not present, the Syt1 dimer, either isolated or within the Syt1-SNARE complex, chelates four Ca^2+^ ions, two by the CBLs of each domain, and no bridging between the C2 domains occurs.

We next asked how the observed formation of the bridged PM-4Ca^2+^Syt1-SNARE-Cpx complex (Fig. 2) would affect the insertion of the C2 domains of Syt1 into PM. The Run 1 was continued for additional 7.2 μs, and the penetration of the C2 domains into PM over the course trajectory was analyzed. We found that CBL-PM insertion for either of the C2 domains did not increase over the course of the trajectory (Fig. 3 A), and for the C2B domain it even slightly diminished. In the end of the trajectory, both C2 domains were tightly attached to PM but not inserted into it (Fig. 3 B, C2 domains bridged).

**Figure 3.**
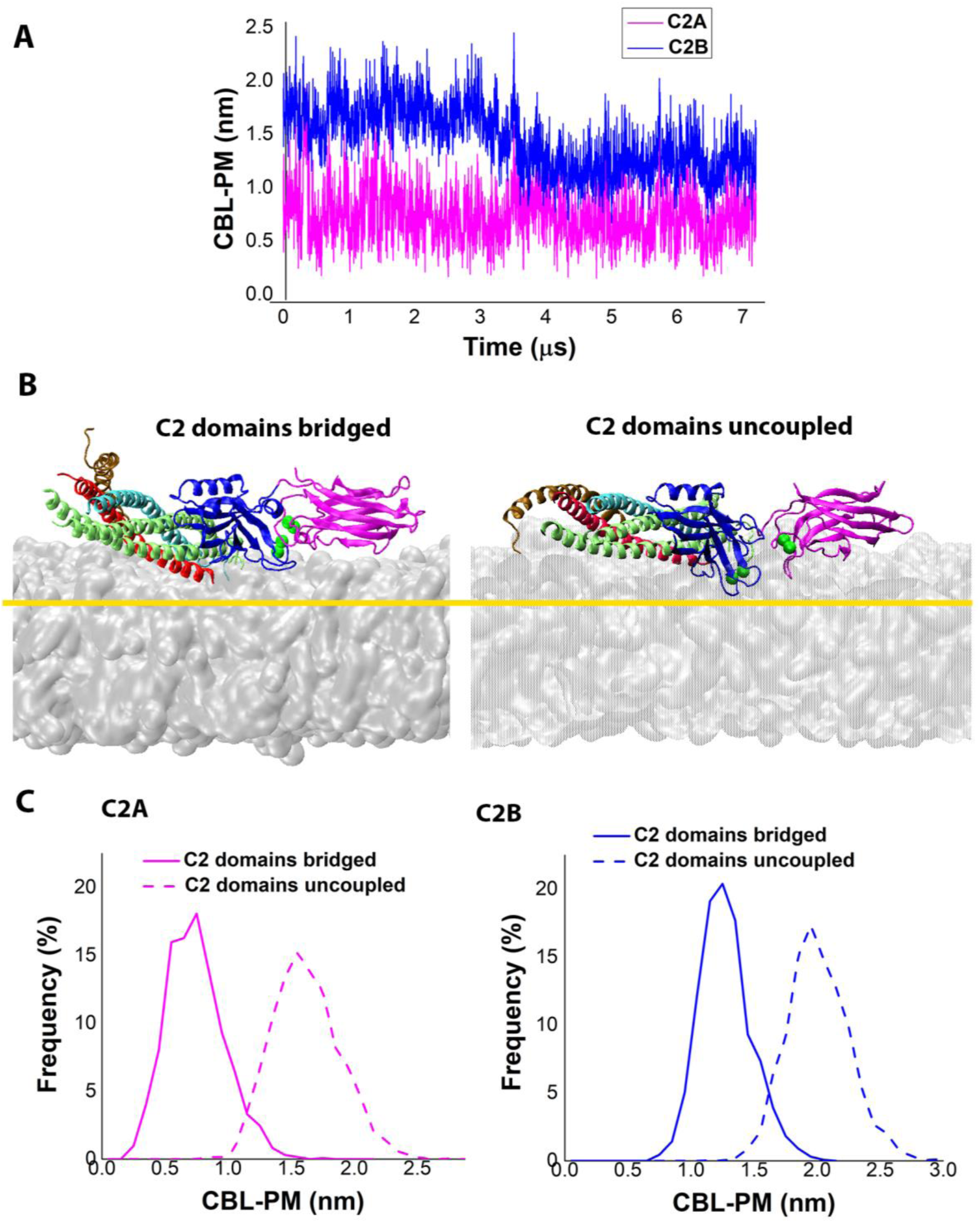
The comparison of the PM-4Ca^2+^Syt1-SNARE-Cpx complexes with bridged and uncoupled C2 domains. **A.** The bridged PM-4Ca^2+^Syt1-SNARE-Cpx complex shows no increase in the penetration into PM over the course of the trajectory Note the steady C2A-PM attachment (magenta) and diminishing C2B-PM penetration (blue). **B.** The bridged (this study) versus uncoupled[2] PM-4Ca^2+^Syt1-SNARE-Cpx complexes in the end of respective trajectories. The yellow line denotes the penetration into PM for the complex with the uncoupled C2 domains. **C.** The complex with the bridged C2 domains has a significantly lower level of the CBL-PM insertion (p<0.001 per K-S test) compared to the complex with the uncoupled C2 domains (replotted from[2]).

This finding contrasts the results of our earlier MD study [2], which investigated the PM-4CA^2+^Syt1-SNARE-Cpx complex with the uncoupled C2 domains, generated by positioning of the 4Ca^2+^Syt1-SNARE-Cpx complex onto PM, followed by a prolonged MD (Fig. 3 B, C2 domains uncoupled). Notably, the 4Ca^2+^Syt1-SNARE-Cpx complex with the C2 domains being uncoupled promotes the insertion of the CBLs into PM (Fig. 3 C), in contrast to the PM-4CA^2+^Syt1-SNARE-Cpx complex with the C2 domains being bridged. This result suggests that bridging of the CBLs by Ca^2+^ ions counteracts the immersion of the CBLs into the lipid bilayer.

Together, the results presented in figure 3 show that the Ca^2+^ bound tips of the C2 domains would insert into PM if Ca^2+^ ions are chelated prior to the Syt1-PM attachment, while the C2 domains would not insert into PM if Ca^2+^ chelation follows the Syt1-PM attachment Syt1 (Fig. 3 B, C). In other words, the PM-4Ca^2+^Syt1-SNARE-Cpx complex can be formed via different pathways, and the specific pathway determines the level of the insertion of the C2 domains of Syt1 into PM.

We questioned therefore what is the sequence of the events and interactions of Syt1 that leads to the formation of the pre-fusion PM-4Ca^2+^Syt1-SNARE-Cpx complex with the C2 domains of Syt1 being able to insert into PM and to drive fusion.

### Syt1 docks to the SNARE bundle via the association of the C2B domain with SNAP25 and PM, and subsequently the PM-Syt1-SNARE-Cpx complex transitions to an intermediate state with the C2A domain of Syt1 attaching to Cpx

We started from simulating the dynamics of full-length Syt1 (Syt1^fl^) attached to SV. The initial 3D structure of human Syt1^fl^ (Fig. 4 A) was generated employing homology modeling as described in Methods. The TM domain (TMD) of Syt1^fl^ was then inserted in the lipid bilayer mimicking SV, and all the lipid monomers overlapping with the TMD of Syt1 were removed (Fig. 4 B). Following the equilibration of the system, a 3.6 µs MD run was performed (Fig. 4, C. D; Video 2). The TMD of Syt1 retained its helical structure over the entire length of the trajectory, remaining inserted into the lipid bilayer. The orientation of the C2A domain remained largely the same over the entire course of the trajectory (Fig. 4 D, magenta), with its CBLs being in the proximity to the lipid bilayer. The linker domain was largely unstructured, with the exception of a short alpha helix formed by the residues 107-118 (DVKDLGKTMKDN), which retained its helical structure over the entire length of the trajectory (Fig. 4 D, black). The linker helix interacted with the lipid bilayer and with the C2A domain, being often positioned between the two (Fig. 4 D: points 1, 3-5). Notably, the C2B domain was highly mobile (Fig. 4 C-E), being typically distanced from the lipid bilayer (Fig. 4 D: points 1, 3, 4, 5, 6, 7), but sometimes attaching to it (Fig. 4 D: points 2, 8).

**Figure 4.**
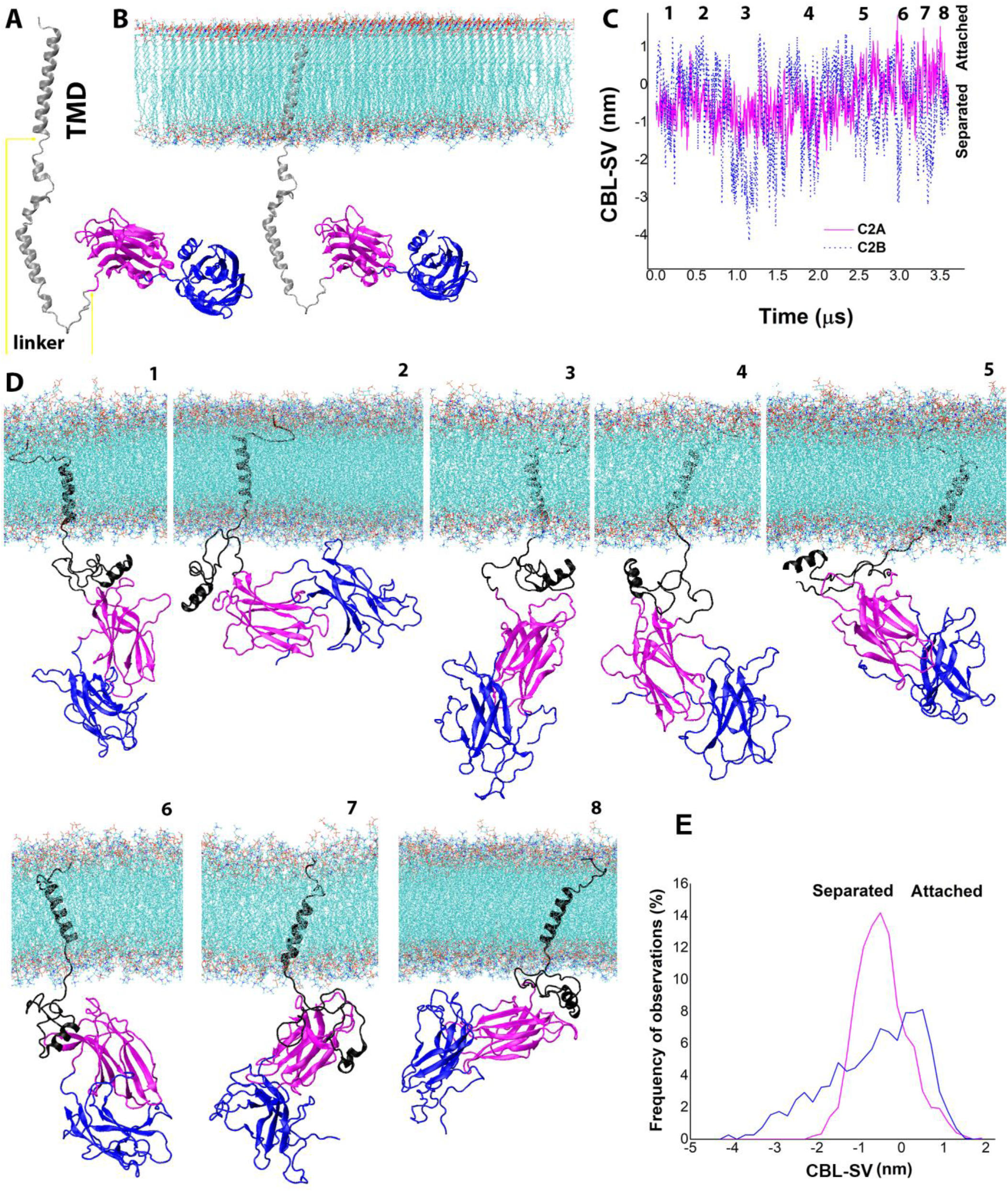
The dynamics of Syt1^fl^ attached to the lipid bilayer mimicking SV. **A.** The initial Syt1^fl^ structure with the linker domain and TMD. **B.** TMD inserted in the lipid bilayer. **C.** The attachment/separation of the CBLs of the C2 domains to/from the SV lipid bilayer over the course of the trajectory. The time-points 1-8 correspond to the states shown in the panel D. **D.** The states of Syt1^fl^-SV complex along the trajectory. **E.** The frequency distributions for the CBL-SV separation/attachment. Note that the C2A domain (magenta) predominantly remains in the SV proximity, while the C2B domain (blue) alternates between distancing and attachment.

The observed mobility of the C2B domain and its distancing from the SV bilayer could be favorable for anchoring to the SNARE bundle or PM, or possibly both. To investigate whether this is the case, we took the final point of the Syt1^fl^-SV trajectory (shown in Fig. 4) and added the SNARE-Cpx complex to the system. The SNARE-Cpx bundle was oriented parallel to the lipid bilayer, so that the C2B domain was facing SNAP25, and Z-dimension of the system was decreased so that the C2B domain approached the leaflet mimicking PM (Fig. 5 A). The system was then equilibrated, and a 6.2 µs MD run was performed.

**Figure 5.**
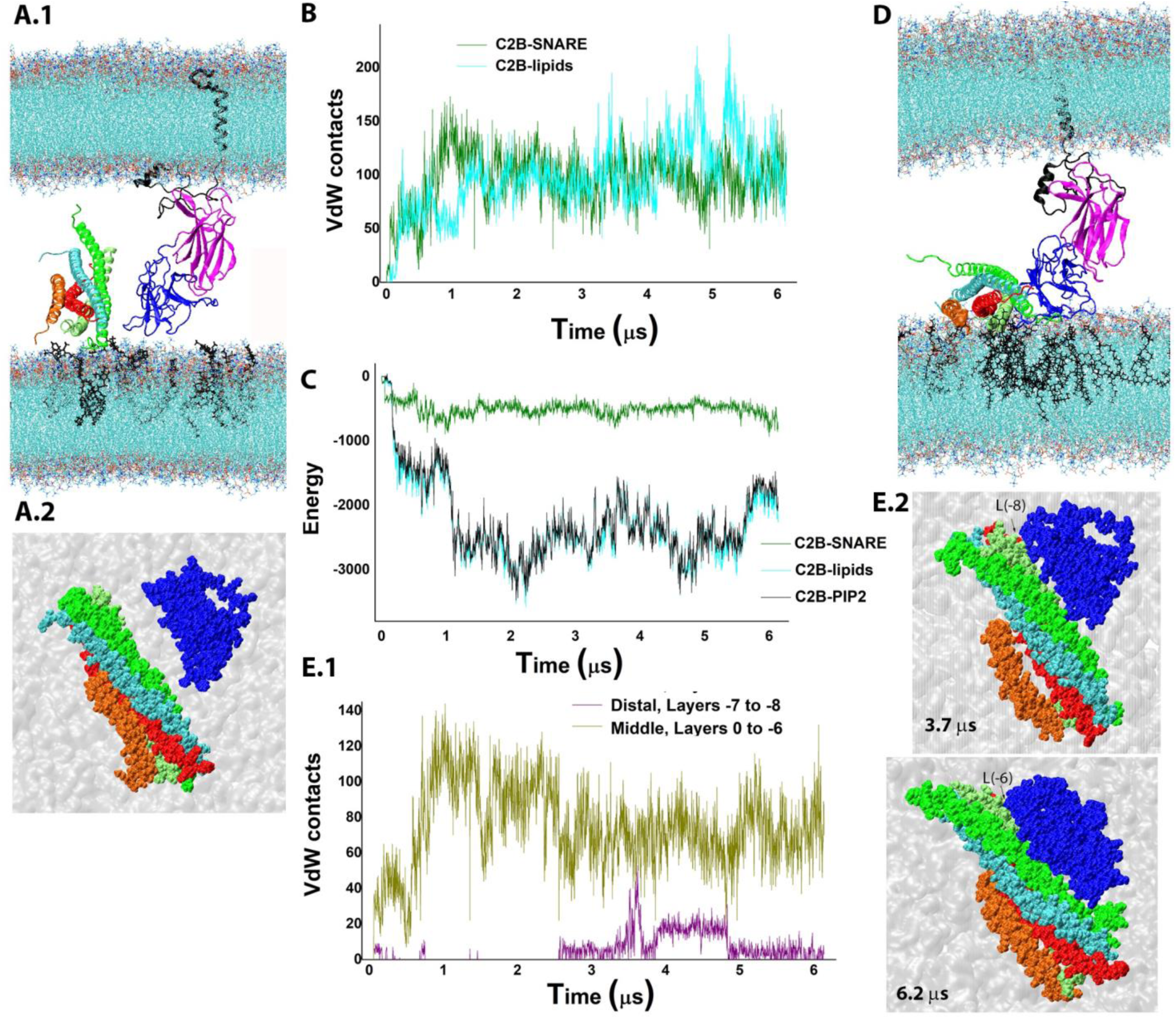
The C2B domain of the SV-attached Syt1^fl^ anchors simultaneously to t-SNARE and PIP_2_. **A.** The initial state of the molecular system. **A.1** The SNARE-Cpx bundle is positioned in the vicinity of the C2B domain (blue) and parallel to the lipid bilayer, with SNAP25 (green) facing Syt1. The height of the cell is adjusted so that the C2B domain is touching the PM bilayer, which incorporates PIP_2_ (black). **A.2.** The top view of the SNARE-Cpx bundle and the C2B domain on PM in the VdW representation. The C2A domain, linker and TMD of Syt1 are removed for clarity. **B.** The number of contacts between the C2B domain and the SNARE bundle (green) rapidly increases, followed by the increase in the number of contacts between the C2B domain and the lipid bilayer mimicking PM (cyan). **C.** The energy of the interactions between the C2B domain and lipids (cyan) dominates over the energy of the C2B-SNARE interactions (green); the former is largely defined by the C2B-PIP_2_ interactions (black). **D.** The final state of the system in the end of the trajectory: the C2B domain is docked tightly to SNAP25 and attached to the PIP_2_ cluster (black). **E.** The C2B domain shifts from the middle part of the bundle to its distal end (N-terminus) and back over the course of the trajectory. **E.1** The number of contacts between the β_1_β_2_ β_5_β_8_ surface of the C2B domain and middle (ochre) or distal (purple) layers of SNAP25. **E.2.** Two snapshots at 3.7 μs (top) and 6.2 μs (bottom) showing the C2B domain in relation to the SNARE-Cpx bundle in VdW representation. The C2A domain, linker and TMD of Syt1 are removed for clarity.

We observed a rapid docking of the C2B domain to the SNARE bundle (Fig. 5 B, green). This was accompanied by and a slower attachment of the C2B domain to the lipid bilayer mimicking PM (Fig. 5 B, cyan). The energy of the C2B-lipid interactions was largely defined by the C2B-PIP_2_ component (Fig. C, black versus cyan), and it dominated over the C2B-SNARE interactions (Fig. 5 C, cyan versus green). This result suggests that the C2B-PM interactions driven by the strong electrostatic attachment of the PB stretch of the C2B domain to PIP_2_ would dominate and largely determine the configuration and dynamics of the Syt1-SNARE docking. The final structure (Fig. 5 D) had the β_1_β_2_ β_5_β_8_ surface of the C2B domain docked to SNAP25 and the PB stretch attached to the PIP_2_ cluster.

Interestingly, over the course of the trajectory we observed the movement of the C2B module along the SNARE bundle (Fig. 5 E). Initially, the C2B module was attached between the ionic layer in the middle of the bundle and the layer L(−6) at its N-terminus. Subsequently, the C2B domain shifted to the distal end of the bundle to the layer L(−8) (Fig. 5 E.1, purple versus ochre; Fig. 5 E.2, top), and finally shifted back more proximally (Fig. 5 E.2, bottom). The observed dynamics are in line with the heterogeneity of the Syt1-SNARE complex [31].

It should be noted however, that our system (Fig. 5) had extra degrees of freedom, since the SNARE bundle was not attached to PM via the TMD of Sx. Therefore, we sought to investigate the dynamics of the PM-Syt1-SNARE-Cpx complex under more realistic conditions, with the TMD of Sx being included and inserted in PM. We substituted Sx within the generated PM-Syt1-SNARE-Cpx complex (Fig. 5 D) by Sx^TM^, which included the TMD of Sx. The model of Sx^TM^ was generated in our earlier study [2]. The zippered domain of Sx within the PM-Syt1-SNARE-Cpx complex was replaced by Sx^TM^ as described in [2]. Subsequently, the TMD of Sx^TM^ was inserted into the lipid bilayer mimicking PM, and the lipid monomers interfering with the TMD were manually removed. To avoid artificial coupling between the TMDs of Sx and Syt1, we removed from the system the TMD of Syt1 together with the linker.

The system was then equilibrated (Fig. 6 A), and the 5.6 µs MD run of the PM-Syt1-SNARE^TM^-Cpx complex was performed (Fig. 6 B, C). Within the initial 300 ns of the simulation, tight contacts were formed between the CBLs of the C2A domain and the region of Cpx (Fig. 6 C, arrow; D). The C2A-Cpx attachment was largely maintained by a salt bridge (D172 of Syt1 – K54 of Cpx) and by strong hydrophobic interactions (F234 of Syt1 – V61 of Cpx), and it remained intact until the end of the 5.6 µs trajectory (Fig. 6 E). Interestingly, a kink between the Cpx central and accessory helixes developed over the course of the trajectory (Fig. 6 B), suggesting that the Syt1-Cpx interactions on the SNARE bundle may affect the Cpx conformation, displacing the Cpx accessory helix.

**Figure 6.**
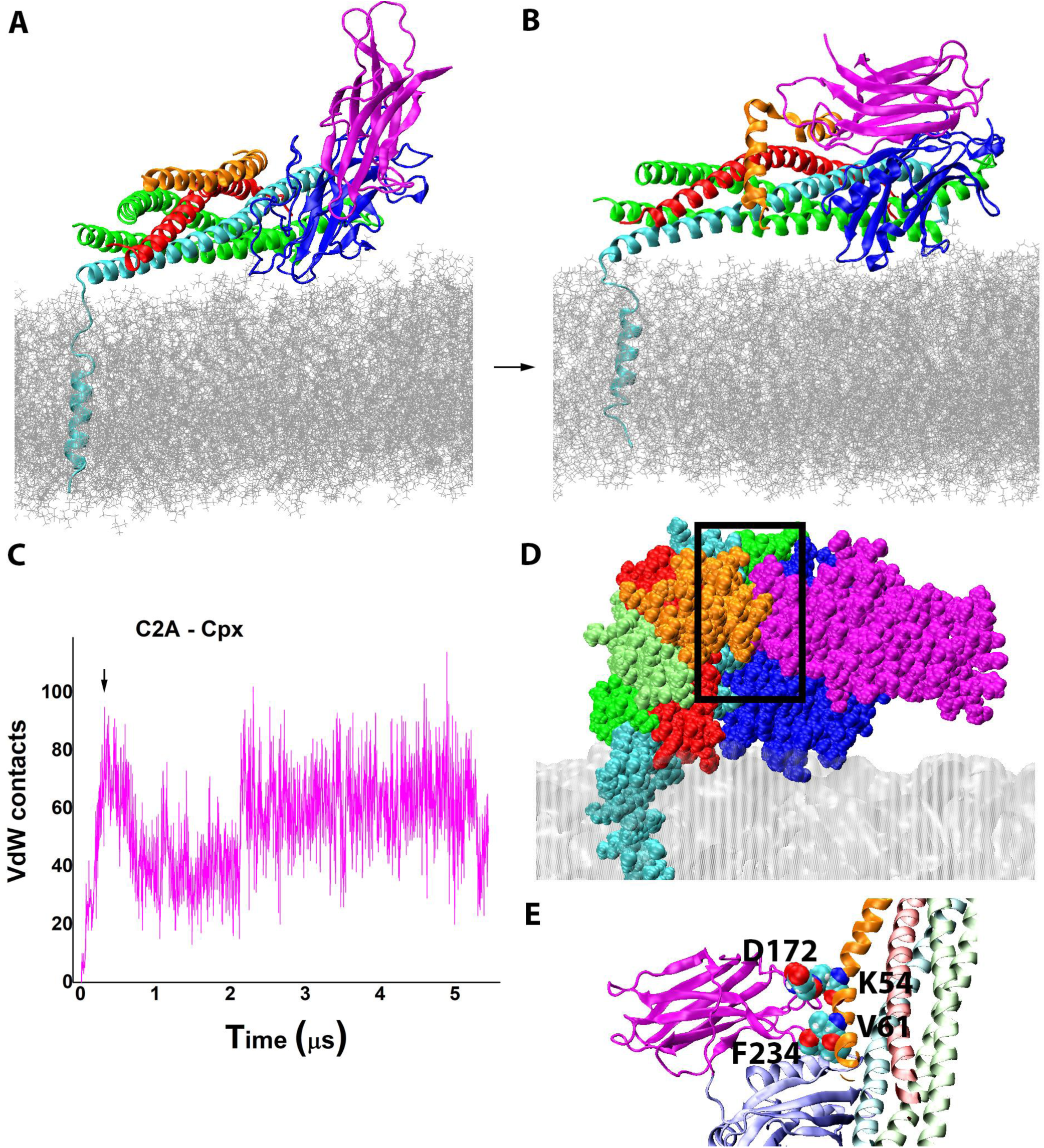
The C2A-Cpx attachment within the PM-Syt-SNARE^TM^-Cpx complex. **A.** The initial state of the complex. Note TMD of Sx (cyan) spanning through PM (silver). **B.** The final state of the complex. Note the CBLs of the C2A domain of Syt1 (magenta) attached to Cpx (orange). **C.** The C2A-Cpx attachment (arrow) is formed within the initial 300 ns of the trajectory and remains intact over the 5.6 µs run. **D.** The VdW representation showing the tight contacts between the CBLs of the C2A domain and Cpx (boxed). **E.** The C2A domain of Syt1 forms a salt bridge (D172-K54) with Cpx, which is strengthened by the hydrophobic interactions (F234-V61).

Thus, our simulations revealed a new conformational state of the PM-Syt1-SNARE^TM^-Cpx complex, which had the CBLs of the C2A domain attached to Cpx in the region connecting the Cpx central and accessory helixes (Fig. 6, B, D, E). Furthermore, our results suggest that SV-attached Syt1 and the PM-attached SNARE-Cpx bundle would likely form this complex when SV and PM are brought into proximity.

### The intermediate PM-Syt1-SNARE^TM^-Cpx complex orchestrates the sequential chelation of Ca^2+^ by the C2B and C2A domains, leading to the formation of the pre-fusion state with both C2 domains of Syt1 inserted into PM

To understand whether the formation of the PM-Syt1-SNARE^TM^-Cpx complex (Fig. 6 B) could lead to the pre-fusion state, we investigated how Syt1 would bind Ca^2+^ within this complex. First, we examined the CBLs of each domain and found that the Ca^2+^-binding pocket of the C2A domain was shielded by Cpx (Fig. 7 A.1). In contrast, the Ca^2+^-binding pocket of the C2B domain was open (Fig. 7 A.2). We concluded therefore that only the C2B domain could chelate Ca^2+^ within thus complex. We then simulated Ca^2+^ binding by the C2B domain employing the method of K^+^/Ca^2+^ replacement[50] as described above. Two Ca^2+^ ions were chelated within the Ca^2+^-binding pocket of the C2B domain (Fig. 7 B), matching the Ca^2+^ coordination obtained in earlier experimental[51] and computational [50] studies.

**Figure 7.**
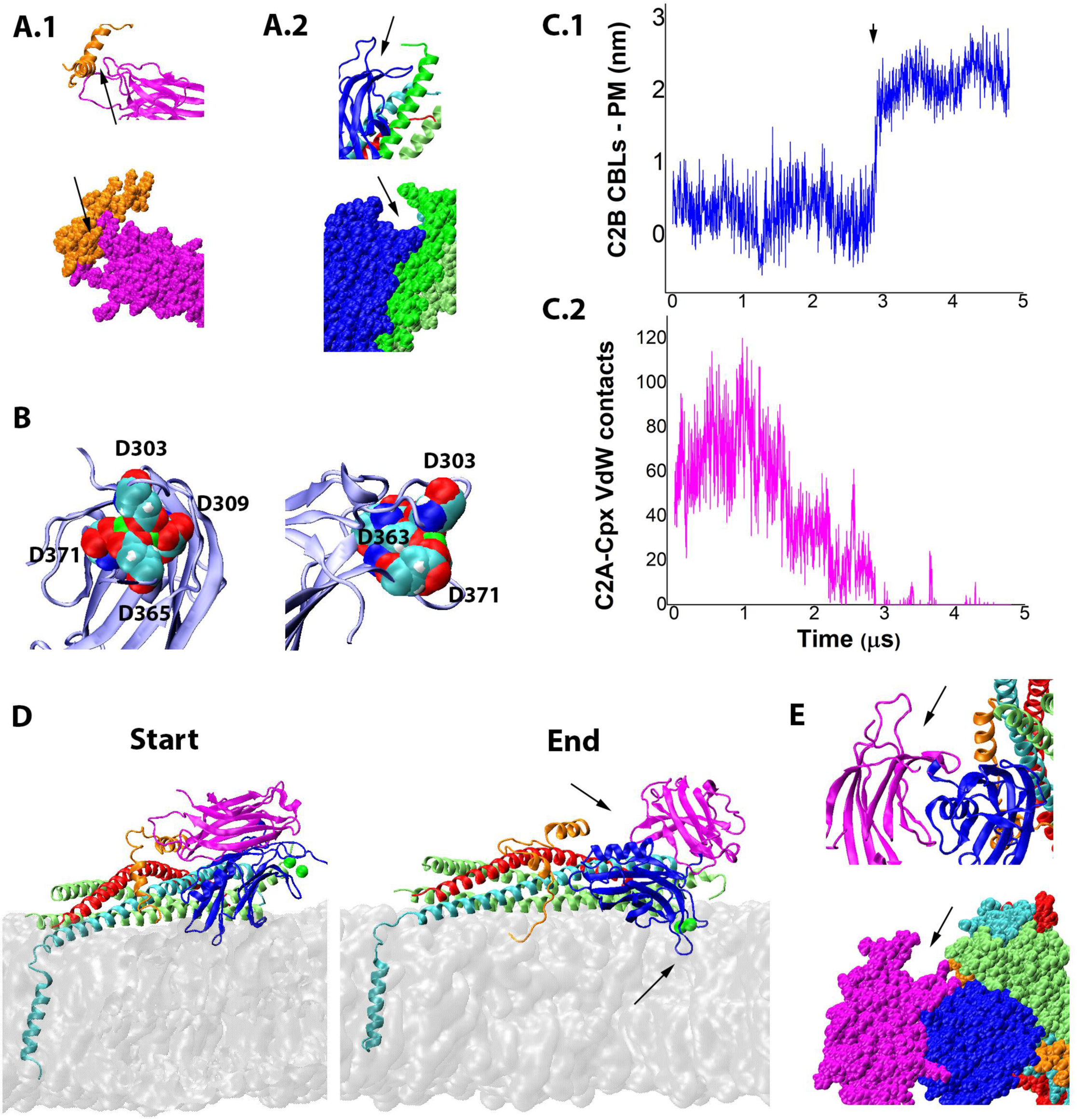
Ca^2+^ binding by the C2B domain drives a conformational transition of the PM-2Ca^2+^Syt1-SNARE^TM^-Cpx complex, so that the tip of the C2B domain imserts into PM and the C2A domain separates from Cpx. **A.** The CBLs of the C2A domains are shielded by Cpx (A.1, arrows; the SNARE bundle is removed for clarity), while the CBLs of the C2B domain are accessible (A.2, arrows). Cartoon (top) and VdW representations (bottom) are shown/ **B.** Two perpendicular views show the aspartic residues (in VdW representation) of the CBLs of the C2B domain binding two Ca^2+^ ions (green). **C.** The trajectory shows a conformational transition (C.1, arrow) leading to the CBLs of the C2B domain inserting into PM, accompanied by the C2A domain separating from Cpx (C.2). **D.** The initial and final states of the PM-2Ca^2+^Syt1-SNARE^TM^-Cpx complex. Note the CBLs of the C2A domain detached from Cpx and the CBLs of the C2B domain immersed into PM (arrows). **E.** The final state of the PM-2Ca^2+^Syt1-SNARE^TM^-Cpx complex has the Ca^2+^-binding pocket of the C2A domain open (arrow). The cartoon (top) and VdW (bottom) representations are shown.

Next, we performed a 4.8 MD run for the obtained PM-2Ca^2+^Syt1-SNARE^TM^-Cpx complex, which had two Ca^2+^ ions bound by the CBLs of the C2B domain (Fig. 7 C, D). Following the initial 3 μs of the simulations, we observed a rapid conformational transition, with the CBLs of the C2B domain immersing into PM (Fig. 7 C.1, arrow). This was accompanied by a disruption of the C2A-Cpx attachment (Fig. 7 C.2, Video 3). Notably, the final state of the PM-2Ca^2+^Syt1-SNARE^TM^-Cpx complex (Fig. 7 D) had the CBLs of the C2A domain open (Fig. 7 E) due to the separation of the C2A domain from Cpx.

This result suggests that the Syt1-Cpx interactions can define the pattern of Ca^2+^ chelation by Syt1. Indeed, only two Ca^2+^ ions can become chelated by the C2B domain within the PM-Syt1-SNARE^TM^-Cpx complex, however a subsequent conformational transition enables Ca^2+^ binding by the C2A domain. To investigate this mechanism further, we simulated the Ca^2+^ chelation by the C2A domain within the generated PM-2Ca^2+^Syt1-SNARE^TM^-Cpx (Fig. 7).

Three Ca^2+^ ions [6] were inserted in the Ca^2+^-binding pocket of the C2A domain by the K^+^/Ca^2+^ replacement method [50], however only two ions remained chelated (Fig. 8 A), while the third Ca^2+^ ion was rapidly lost from the pocket. This finding is in line with earlier experimental [43, 52] and computational [50] studies, which suggested that the C2AB tandem can only chelate four Ca^2+^ ions, two within each of the C2 domains. To investigate the dynamics of the obtained PM-4Ca^2+^Syt1-SNARE^TM^-Cpx complex, we performed a prolonged (26.5 μs) MD run (Fig. 8, B-D). The initial 12 μs of the trajectory were characterized a relative stability of the complex, with the C2A domain remaining separated from PM (Fig. 8B), although a transient shift of the C2B domain along the bundle (Fig. 8 C) was observed. Subsequently, a conformational transition took place (Fig. 8 B, arrows), so that the C2A domain moved and rotated with its CBLs attaching to PM (Video 4). This was accompanied by a brief detachment of the C2B domain from the SNARE complex (Fig. 8C, arrows) and a subsequent shift of the C2B domain towards the middle of the bundle (Fig. 8C, note the lack of interactions with the distal layers in the end of the trajectory). Following 21 μs of the simulations, the CBLs of the C2A domain immersed into PM (Fig. 8 B, arrow). Importantly, the end-point of the trajectory had the CBLs of both C2 domains inserted into PM (Fig. 8 D), in contrast to the “dead-end” state of the complex (Fig. 8 E).

**Figure 8.**
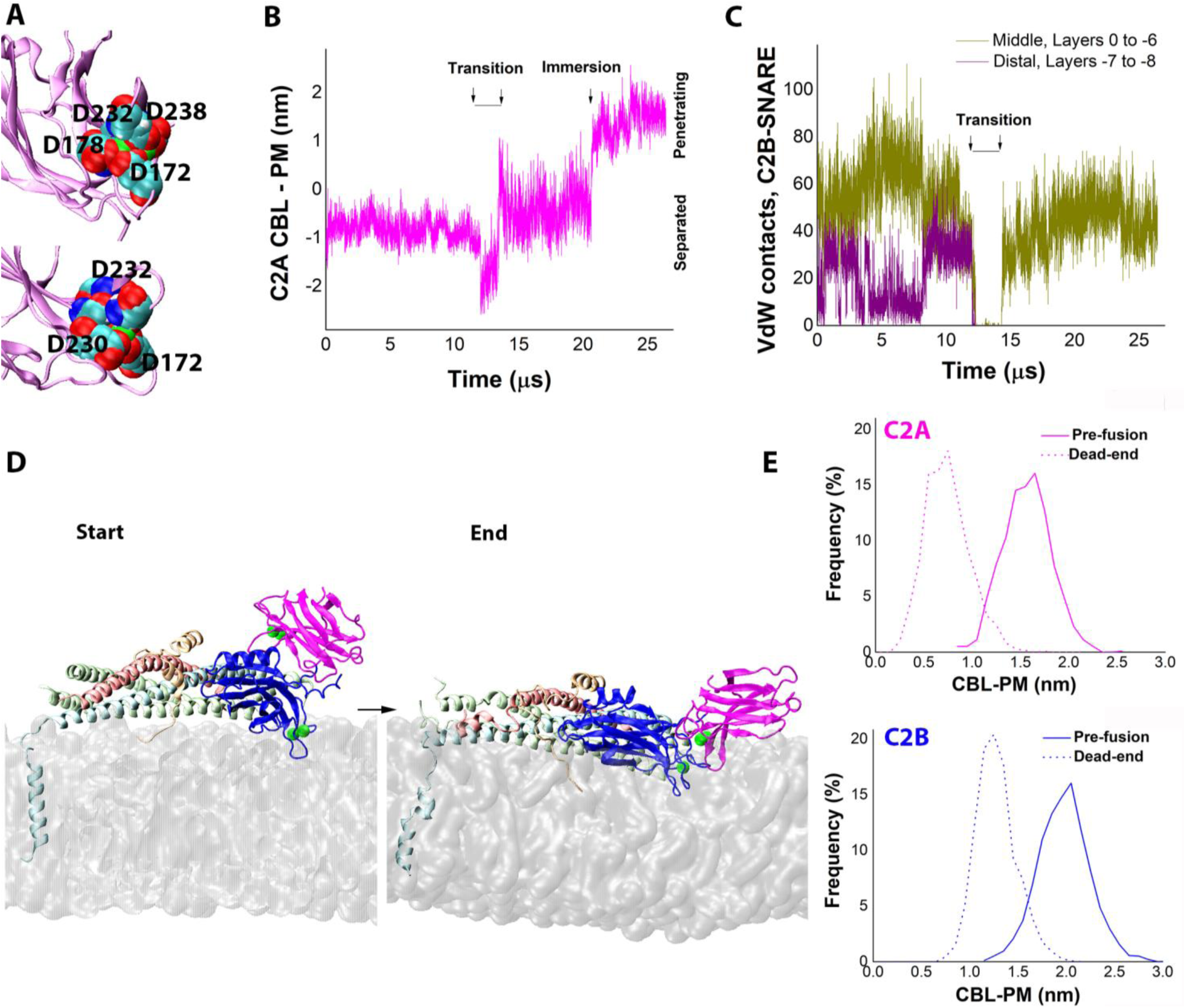
The PM-4Ca^2+^Syt1-SNARE^TM^-Cpx complex transitions to its pre-fusion state with the tips of both C2 domains penetrating into PM. **A.** The CBLs of the C2A domain chelate two Ca^2+^ ions (green), which from coordination bonds with five aspartic acids (shown in VdW representation). Two perpendicular views are shown. **B.** The CBLs of the C2A domain are separated from PM at the initial part of the trajectory (note negative CBL-PM), while a subsequent conformational transition (arrows) brings the CBLs in contact with PM (note CBL-PM fluctuating around 0), which is followed by the insertion of the CBLs into PM (arrow, note increasing CBL-PM). **C.** The C2B domain shifts along the SNARE bundle, interacting with its distal layers (purple) in the beginning of the trajectory, but attaching to the middle of the bundle (ochre) after the conformational transition (arrows). **D.** The initial and final states of the trajectory. Note that in the end of the trajectory the CBLs of both domains with chelated Ca^2+^ ions penetrate into PM. **E**. The levels of the immersion of the C2 domains into PM versus the dead-end state (p<0.001 per K-S test, the dead-end state is replotted from Fig. 3 C). The frequency distributions were constructed for the final 1.5 μs of the trajectory.

Thus, our MD simulations (Fig. 5-8) produced the dynamic model for the formation of the prefusion PM-4Ca^2+^Syt1-SNARE^TM^-Cpx complex. The pre-fusion complex is characterized by the tips of the C2 domains of Syt1 penetrating into PM. The simulations suggested that this pathway is guided by the direct interactions between Syt1 and Cpx on the SNARE bundle (Fig. 6), and that these interactions define the pattern of Ca^2+^ chelation by Syt1 (Fig. 7, 8).

## Discussion

We employed MD simulations to elucidate the conformational pathway that leads to the formation of the pre-fusion protein-lipid complex (Fig. 9). Our results suggest that the C2B domain of Syt1 (Fig. 9 (1)) docks to SNAP25 via its β_1_β_2_ β_5_β_8_ surface, and almost simultaneously the C2 domain attaches to PM via the formation of the salt bridges between its PB stretch and PIP_2_ (Fig. 9 (2)). Subsequently, the CBLs of the C2A domain of Syt1 bind Cpx via ionic and hydrophobic interactions, and the complex transitions to an intermediate state (Fig. 9 (3)). This intermediate state allows the Ca^2+^ chelation by the C2B domain but not by the C2A domain. Binding Ca^2+^ by the C2B domain triggers a conformational switch, so that the C2B domain tilts with its CBLs immersing into PM, and consequentially the C2A domain detaches from Cpx, opening the Ca^2+^-binding pocket (Fig. 9 (4)). The subsequent Ca^2+^ chelation by the C2A domain leads to the transitioning into the pre-fusion state, which is characterized by the tips of both C2 domains being inserted into PM (Fig. 9 (5)). Our simulations also suggest that if this pathway is disrupted, the protein-lipid complex can transition into the “dead-end” state (Fig. 9, 1->6->7), which has both C2 domains of Syt1 tightly attached to PM but not penetrating into it.

**Figure 9.**
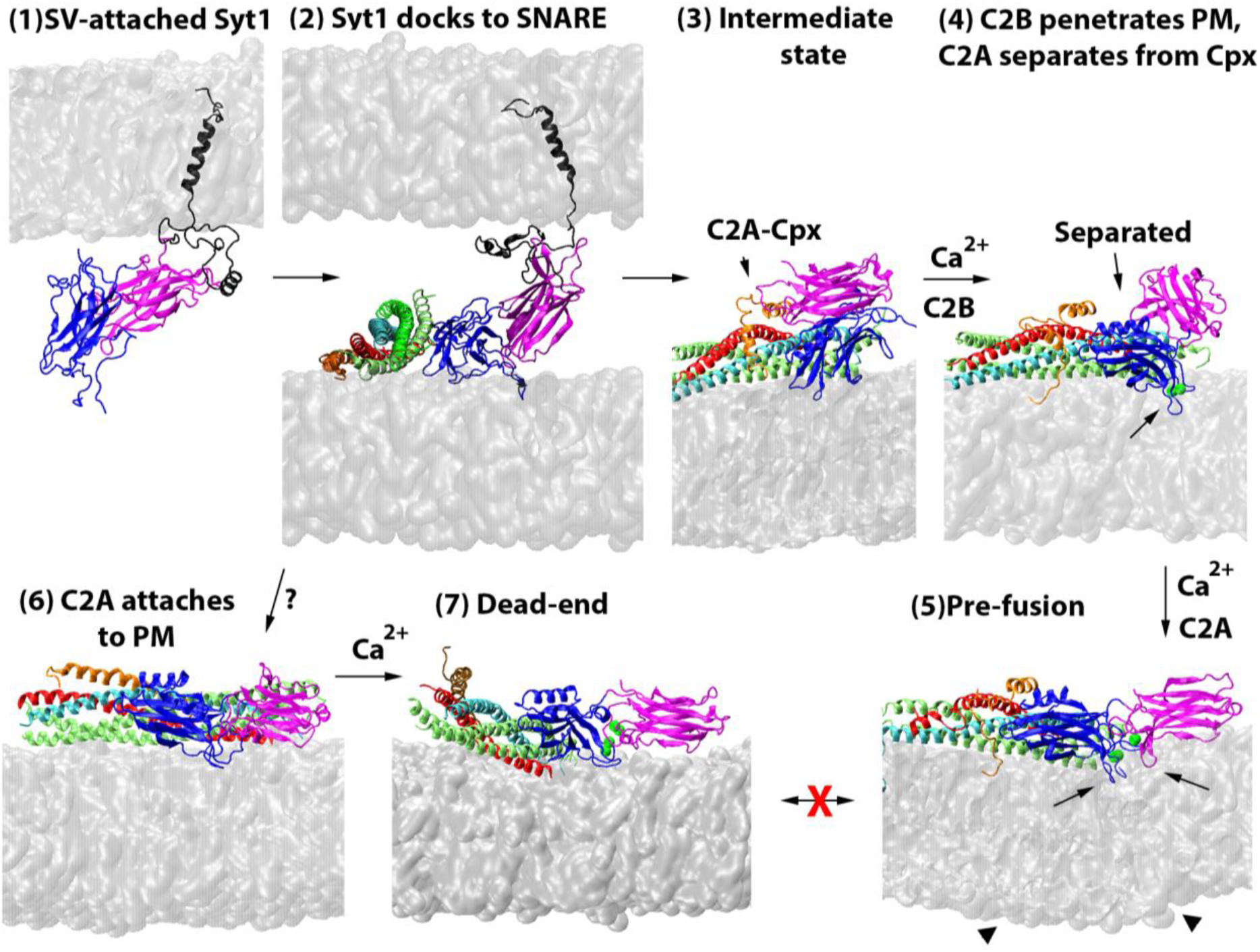
The model of the dynamic formation of the prefusion complex. The SV-attached Syt1 (1) docks to the SNARE bundle and to PM via its C2B domain (2). Subsequently, the intermediate state of the PM-Syt1-SNARE-Cpx complex is formed via the attachment of the C2A domain of Syt1 to Cpx (3). Ca^2+^ chelation triggers the immersion of the CBLs of the C2B domain into PM followed by the separation of the C2A domain from Cpx (4; arrows). The subsequent Ca^2+^ chelation by the C2A domain leads to the pre-fusion state (5) which has the tips of both C2 domains penetrating into PM (arrows), which in turn promotes PM curvature (arrowheads). An alternative pathway may involve the attachment of both C2 domains to PM prior to the Ca^2+^ chelation (6). The subsequent Ca^2+^ binding would lead to the dead-end state (7), with the C2 domains of Syt1 being bridged and tightly attached to PM but not penetrating into the lipid bilayer.

### Coordinated insertion of the tips of the C2 domains of Syt1 into PM is a critical step in synaptic fusion, and it depends on the interactions of Syt1 with the SNARE complex

Biophysical studies of membrane fusion suggested that Syt1 triggers exocytosis via the insertion of the tips of its C2 domains into PM, locally promoting PM curvature, which in turn triggers fusion[3, 9, 53]. Further evidence for this model came from a demonstrated correlation between membrane curvature and the rate of exocytosis[54]. Notably, optical studies showed that the C2A and C2B domains act synergistically, interacting in the insertion into PM and in membrane bending [55]. Thus, it is generally agreed that the synergistic coordinated insertion of the tips of the C2 domains of Syt1 into PM is a critical step in synaptic fusion.

Syt1 interacts with the SNARE complex [56], and these interactions, largely mediated by SNAP25, are essential for exocytosis [15, 57, 58]. Studies at PC12 cells demonstrated that Syt1 utilizes SNARE binding to promote Ca^2+^-dependent membrane bending and the fusion pore expansion [14]. Furthermore, our earlier MD study demonstrated that the interaction of Syt1 with the SNARE complex is required to uncouple the C2 domains of Syt1 and to promote their insertion into PM[2]. Together, these studies suggest that the insertion of the C2 domains of Syt1 into PM depends on the Syt1-SNARE attachment.

The present study shows that the penetration of the Ca^2+^-bound tips of the C2 domains of Syt1 into PM requires a sequence of Syt1 interactions and conformational transitions. We outline the conformational pathway that can lead to the formation of the pre-fusion PM-4Ca^2+^Syt1-SNARE-Cpx complex, as well as an alternative “dead-end pathway”.

### The Syt1-SNARE-Cpx complex can adopt multiple conformational states enroute to triggering fusion

The Syt1-SNARE complex was shown to adopt multiple conformational states *in vitro* [1, 18, 30–32] being structurally heterogenous. These studies raised the question of the structure and dynamics of this complex *in vivo,* between the lipid bilayers of SV and PM. Importantly, the interactions of Syt1 with lipids depend on the lipid composition[59], and in particular on the PM component PIP_2_ [60, 61]. Therefore, the proximity to PM could define the structure of the Syt1-SNARE complex.

The MD approach enabled building all-atom models of the Syt1-SNARE-Cpx complexes interacting with PM [2, 33]. The models relied on the large primary interface between the ***β_1_β_2_ β_5_β_8_*** surface of the C2B domain and t-SNARE, which was identified by crystallography [18], and validated by a mutagenesis study in *Drosophila*[17]. Notably, MD simulations showed that the formation of the pre-fusion PM-4Ca^2+^Syt1-SNARE-Cpx complex promotes the immersion of both C2 domains of Syt1 into PM and drives the SV-PM merging [2].

Here, we investigated the dynamic formation of the pre-fusion protein-lipid complex. We identified the sequence of the interactions and conformational transitions of Syt1 (Fig. 9, 2-5), starting from docking of the C2B domain to SNAP25 and PM, proceeding via the formation of the intermediate PM-Syt1-SNARE-Cpx complex characterized by the direct Syt1-Cpx attachment, undergoing a two stage Ca^2+^ chelation, and finally inserting the tips of the C2 domains into PM. Our study suggests that the structural heterogeneity of the Syt1-SNARE complex observed *in vitro* reflects the dynamic flexibility of this complex required for its *in vivo* function.

### Cpx may accelerate evoked release by promoting the intermediate state that leads to the formation of the pre-fusion protein-lipid complex

Our MD simulations unraveled the conformational pathway (Fig. 9, 2-5) that requires a direct role of Cpx (Fig. 9 (3)). Cpx was shown to ubiquitously promote Ca^2+^-dependent evoked transmission[24] and also to clamp spontaneous or asynchronous activity at several preparations[62]. Although the Cpx clamping function is largely understood, being mediated by its accessory helix [63–66], the molecular mechanisms of the Cpx facilitatory action in evoked release are still debated[24, 62, 67, 68]. However, structural[30, 32], genetic[25] and functional[27, 28] studies suggest that Cpx may regulate evoked release by directly binding Syt1.

Our MD simulations led to the all-atom model of the intermediate PM-Syt1-SNARE-Cpx complex (Fig. 9 (3)) which has the C2A domain of Syt1 directly binding Cpx. Within this complex, the CBLs of the C2A domain attach to the region of Cpx that connects its central and accessory helixes, forming a salt bridge supported by strong hydrophobic interactions. Notably the Syt1-binding region of Cpx is positioned in the vicinity of the switch-breaker mutation [66] which was shown to selectively inhibit evoked transmission. Our MD simulations show that the Syt1-Cpx attachment affects both proteins by 1) displacing the accessory helix of Cpx from the SNARE bundle, which could diminish the Cpx clamping function[64, 65], and 2) blocking the Ca^2+^ binding pocket of the C2A domain of Syt1. The latter mechanism enables Cpx to orchestrate the sequential Ca^2+^ chelation by the C2B and C2A domains and to prevent the formation of the “dead-end” state of the PM-Syt1-SNARE-Cpx complex.

### A conformational switch defines the pattern of Ca^2+^ chelation by Syt1

SVs dock to PM prior to releasing transmitters in response to a Ca^2+^ inflow, and synchronous release is predominantly triggered by Ca^2+^ binding to the Syt1 molecules of SVs that are docked in the vicinity of Ca^2+^ channels[11–13, 69]. The affinity of Syt1 to Ca^2+^ is relatively low, however it increases remarkably in the presence of the PM component PIP_2_ [52, 70, 71]. Therefore, a cooperative action of Ca^2+^ and PIP_2_ has been proposed[72]. In line with these studies, our MD simulations suggest that the C2B domain of Syt1 chelates Ca^2+^ after becoming attached to PM via the salt bridges between its PB stretch and PIP_2_ (Fig. 9 (3)).

Our simulations also show that the C2A module, in contrast, should bind Ca^2+^ prior to its attachment to PM, since otherwise the C2A and C2B domains bridge their CBLs, and this counteracts their insertion into PM (Fig. 9 (6)). We show that the sequential Ca^2+^ chelation by the C2B and then the C2A domains can be orchestrated by a fast Cpx-dependent conformational switch. This switch acts at a sub-microsecond scale and releases the Cpx block from the C2A domain upon Ca^2+^ chelation by the C2B domain (Fig. 9, 3->4). This model agrees with earlier studies suggesting that the C2 domains of Syt1 are not equivalent in triggering fusion, and that the C2B domain has the primary role [73–75].

### Promoting the dead-end state could inhibit fusion

Our simulations suggest that the Cpx-mediated conformational switch prevents the “dead-end” scenario, which has the PM-4Ca^2+^Syt1-SNARE-Cpx complex formed but not distorting PM (Fig. 9 (7)). Therefore, it is likely that the protein-lipid complex can transition into its dead-end state in the absence of Cpx. This would account for the reduced and desynchronized evoked release in the Cpx knockout preparations [20, 23, 25, 63].

Importantly, the mutations in Syt1 favoring the “dead-end” state could become dominant-negative, since the accumulation of the dead-end complexes could block the available release sites and preclude native Syt1 forms from docking to t-SNARE clusters and PIP_2_-rich PM patches. Notably, it was demonstrated that neutralizing Asp residues of the CBLs of the C2B domain of Syt1 produces the dominant negative phenotype in *Drosophila* [17, 73]. Furthermore, several disease-associated missense mutations in the CBLs of the C2B domain have been discovered, all causing developmental neuropathies as heterozygotes [76–78]. These mutations alter the attachment of the CBLs of the C2B domain to PM[79], and hypothetically they could drive the transition of the protein-lipid complex into its “dead-end” state (Fig. 9 (6,7)). Further modeling and experimentation will be needed to test this hypothesis.

## Acknowledgements

Anton2 computer time was provided by the Pittsburgh Supercomputing Center (PSC) through Grant R01GM116961 from the National Institutes of Health. The Anton2 machine at PSC was generously made available by D.E. Shaw Research. This work used Anvil and Stampede supercomputers through allocation from the ACCESS program, which is supported by National Science Foundation grants #2138259, #2138286, #2138307, #2137603, and #2138296.

